# Frequency of mispackaging of *Prochlorococcus* DNA by cyanophage

**DOI:** 10.1101/2020.02.18.953059

**Authors:** Raphaël Laurenceau, Nicolas Raho, Mathieu Forget, Aldo Arellano, Sallie W. Chisholm

## Abstract

*Prochlorococcus* cells are the numerically dominant phototrophs in the open ocean. Cyanophages that infect them are thus a notable fraction of the total viral population in the euphotic zone, and, as vehicles of horizontal gene transfer, appear to drive their evolution. Here we examine the propensity of three cyanophages – a podovirus, a siphovirus, and a myovirus – to mispackage host DNA in their capsids while infecting *Prochlorococcus,* the first step in phage-mediated horizontal gene transfer. We find the mispackaging frequencies are distinctly different among the three phages. Myoviruses mispackage host DNA at low and stable frequencies, while podo- and siphoviruses vary in their mispackaging frequencies by orders of magnitude depending on growth light intensity. We attribute this difference to the concentration of intracellular reactive oxygen species and protein synthesis rates. Based on our findings, we propose a model of mispackaging frequency determined by the imbalance between the production of capsids and the number of phage genome copies during infection.

## INTRODUCTION

With an estimated population of 3 × 10^27^ cells on Earth, *Prochlorococcus* is the most numerically abundant phytoplankton species in the global oceans [1]. Distributed throughout the euphotic zone in mid-latitude habitats, *Prochlorococcus* is responsible for fixing an estimated 4 billion tons of carbon each year, playing a central role in ocean food webs. *Prochlorococcus* genomes are highly streamlined, an adaptation for growth in oligotrophic surface waters [2]. Different ocean regions contain hundreds to thousands of *Prochlorococcus* coexisting subpopulations [3], [4]. Horizontal gene transfer has had a large impact on *Prochlorococcus* evolution, both in terms of generating genomic variability in genomic islands [5] or in reinforcing genetic similarity in closely related cells [6], [7]. Furthermore, ways of exchanging genetic information among cells are particularly critical for streamlined organisms such as *Prochlorococcus*; its absence could lead to extinction in the face of environmental change [8], [9].

Modes of horizontal gene transfer (HGT) available to *Prochlorococcus* are lipid-bound vesicles, which contain DNA and are known to be abundant in the oceans [10], [11], natural transformation, but only in some LLIV clades that contain the necessary genes for competence [12], and transduction via cyanophage capsids. The latter is one of the canonical modes of HGT in bacteria [13]-[15] and occurs at significant frequencies in aquatic ecosystems [16]-[18]. Given the abundance of cyanophages that infect *Prochlorococcus* in the oceans (easily reaching 10^5^ to 10^6^ phages ml^-1^) [19], [20], which represent a notable fraction of the total viral population in the euphotic zone [21], it seems self-evident that transduction must be occurring. Thus, we focused this study on exploring and quantifying this phenomenon.

Specialized (reviewed in [22], [23]) and lateral [24] transduction in bacteria are strictly mediated by lysogenic phages, as they enter and leave their host chromosome. Prophages, the hallmarks of lysogeny, however, have not been observed in the hundreds of *Prochlorococcus* isolates and single-cell genomes from the wild [25], suggesting that the vast majority of phages infecting *Prochlorococcus* are strictly lytic. This leaves generalized transduction - the random mispackaging of host DNA into capsids and its subsequent injection and recombination in a recipient cell - as the dominant route of phage-mediated HGT in these bacteria.

To explore the potential of *Prochlorococcus* lytic cyanophages to be vectors of generalized transduction, we quantified the mispackaging of fragments of host DNA during infection of *Prochlorococcus* MED4 by three cyanophages with different morphotypes belonging to the sipho- (P-HS2), myo- (P-HM2) and podovirus (P-SSP7) families (Fig. 1A). Our goal was to determine the frequency of mispackaging in the different morphotypes and begin to explore the environmental factors that might influence mispackaging events in the wild.

**Fig. 1.**
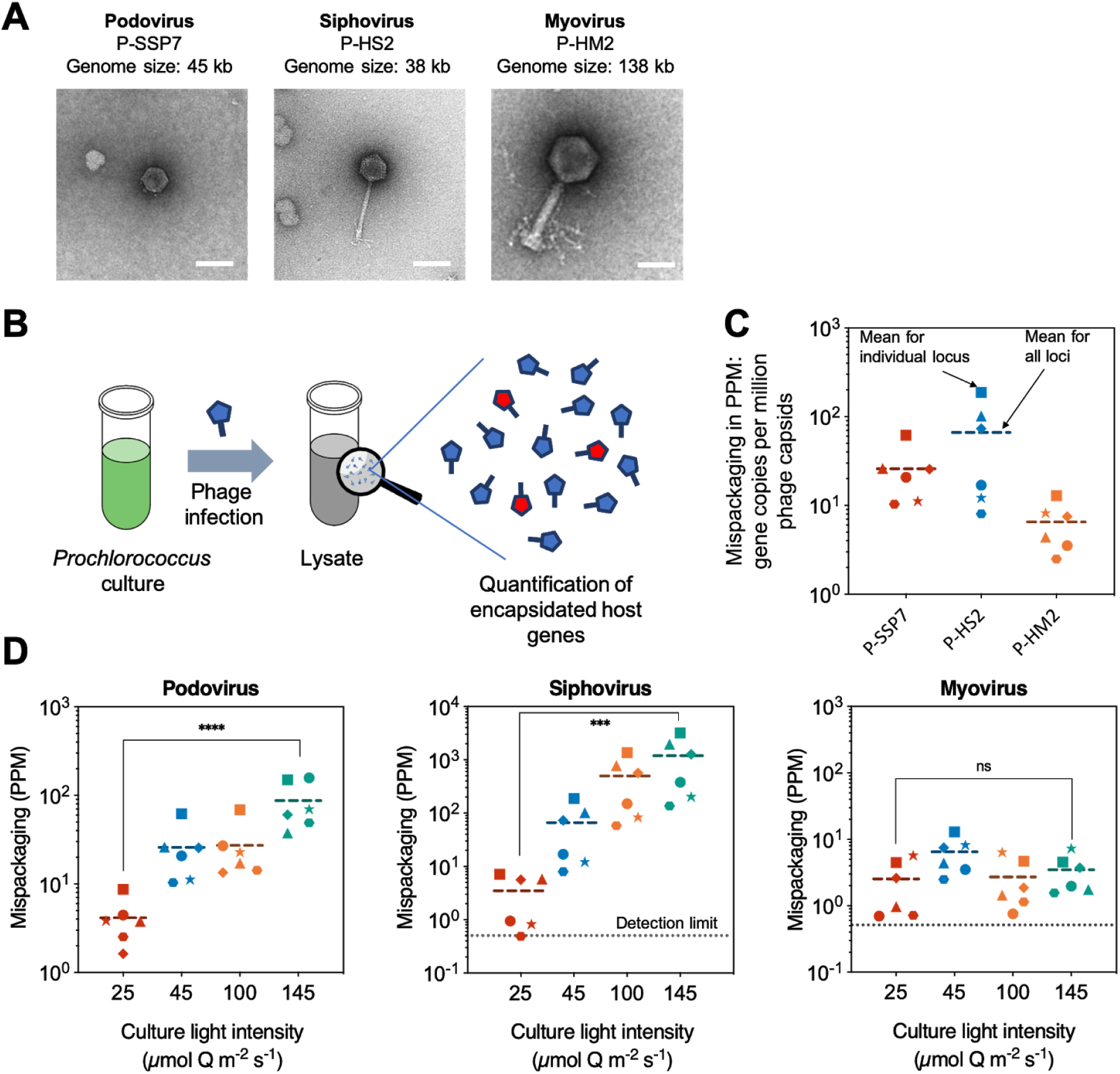
**A)** Electron micrograph of the three cyanophages infecting *Prochlorococcus* MED4 used in this study. The scale bar is 50 nm. **B)** Cartoon illustrating the experimental method. A cyanophage infection typically results in the production of a small fraction of cyanophage capsids having mispackaged host DNA, represented in red. **C)** Detection of host DNA mispackaging inside cyanophage capsids under standard laboratory conditions, at a constant light intensity of 45 μmol Q m^-2^ s^-1^ (see methods for details). The frequency of a gene mispackaging during infection is expressed in PPM: gene copies per million cyanophage capsids. Each symbol is the mean of three parallel infections for a given locus (Table 3). The colored dotted line is the mean value of all loci and all replicates. The 3 colors for the different phage are arbitrary – for ease of visualization. **D)** Impact of growth light intensity on mispackaging during infection. The data 45 μmol Q m^-2^ s^-1^ are from panel C, replotted for comparison. The dotted line at 0.5 PPM, marks the detection limit of the assay. The 4 colors for the different phage are arbitrary – for ease of visualization.

## RESULTS AND DISCUSSION

### Host DNA mispackaging frequency in cyanophage capsids

Using six different loci on the *Prochlorococcus* MED4 chromosome (representative of the entire chromosome, see Table 3), we quantified the level of mispackaging of MED4 DNA in the capsids of the three different cyanophages (Fig. 1A, Table 2) during infection (Fig. 1B, C). The packaging strategy of the podovirus is T7-like [26], the myovirus’ is likely T4-like headful packaging [27] while the siphovirus’ is unknown, but 490 bp direct terminal repeats suggest a T7-like packaging mechanism [28]. Host DNA was detected in the capsids of all 3 cyanophages (Fig. 1C). The levels of mispackaging are low compared to those reported for high transducing phages in *S’, aureus* [29] for example, but comparable to levels (~10 PPM) reported for the marine *Synechococcus* cyanophage S-PM2 [30].

**Table 1.** Cyanophage growth parameters in response to host cell growth light intensity, measured by qPCR of phage genome copies in the extracellular and intracellular fraction. The burst size and latent period were calculated for each replicate infection independently, and results correspond to the arithmetic mean of triplicate infection with standard errors. Low light: 25 μmol Q m^-2^ s^-1^, High light: 145 μmol Q m^-2^ s^-1^.

| Cyanophage | Podovirus<br>P-SSP7 |  | Siphovirus<br>P-HS2 |  | Myovirus<br>P-HM2 |  |
| --- | --- | --- | --- | --- | --- | --- |
| Light intensity<br>(acclimated<br>host cells) | Low<br>light | High<br>light | Low<br>light | High<br>light | Low<br>light | High<br>light |
| Burst Size | 55 ± 7 | 20 ± 6 | 40 ± 6 | 146 ± 40 | 5 ± 2 | 5 ± 1 |
| Latent Period<br>(hours) | 24 ± 2 | 20 ± 2 | 23 ± 2 | 17 ± 5 | 8 | 8 |

**Table 2.**
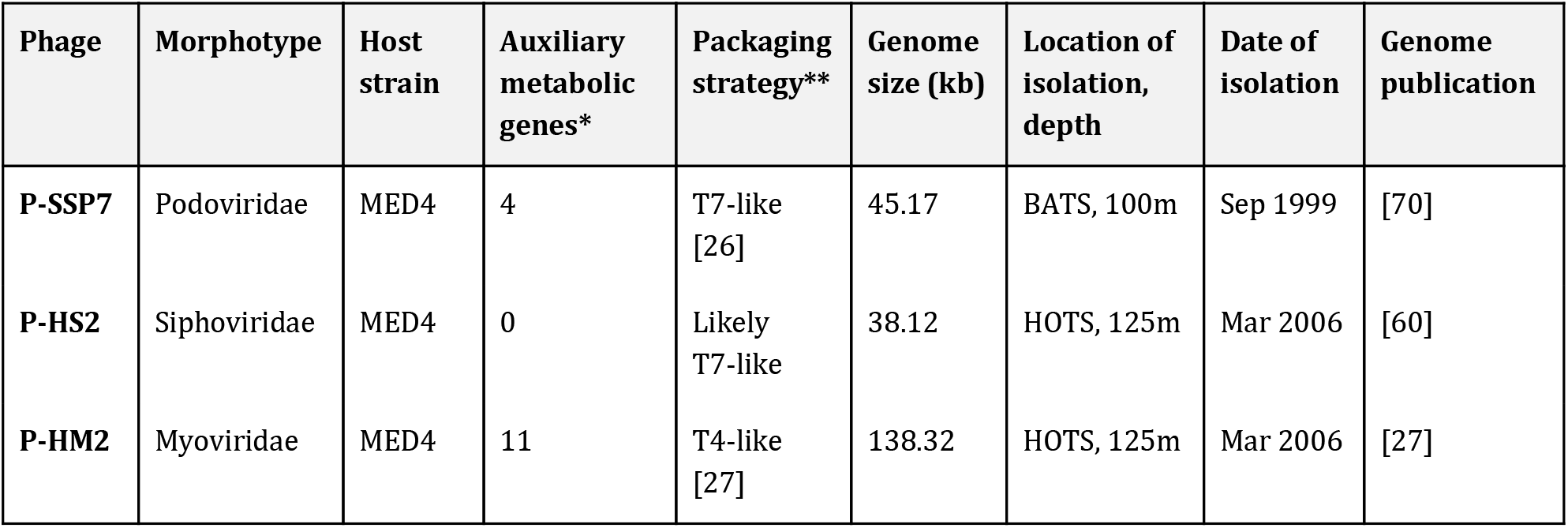
Cyanophages used in this study. HOTS: Hawaii Ocean Time Series station, Pacific oligotrophic gyre. BATS: Bermuda Atlantic Time Series station, Sargasso Sea. (*) Number of different auxiliary metabolic genes related to either carbon metabolism, photosynthesis, DNA synthesis or nutrient uptake processes based on references [37], [60]. (**) Packaging strategy classification based on reference [28].

| Phage | Morphotype | Host strain | Auxiliary metabolic genes* | Packaging strategy** | Genome size (kb) | Location of isolation, depth | Date of isolation | Genome publication |
| --- | --- | --- | --- | --- | --- | --- | --- | --- |
| P-SSP7 | Podoviridae | MED4 | 4 | T7-like [26] | 45.17 | BATS, 100m | Sep 1999 | [70] |
| P-HS2 | Siphoviridae | MED4 | 0 | Likely T7-like | 38.12 | HOTS, 125m | Mar 2006 | [60] |
| P-HM2 | Myoviridae | MED4 | 11 | T4-like [27] | 138.32 | HOTS, 125m | Mar 2006 | [27] |

**Table 3.**
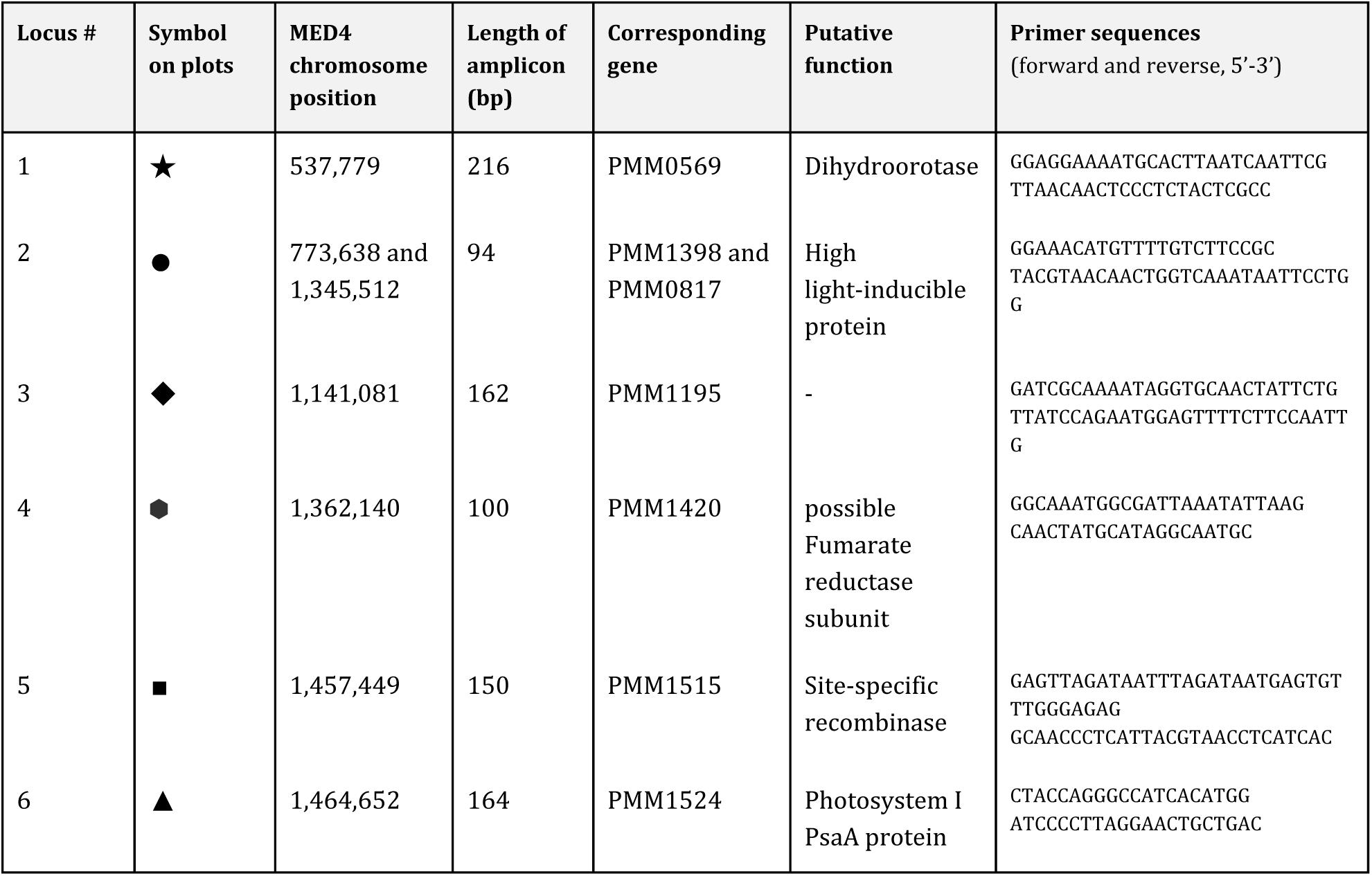
List of *Prochlorococcus* MED4 loci used for qPCR quantification of host DNA inside capsids. Various loci were initially picked to look for specific gene mispackaging. Overall, our results showed a homogeneous mispackaging across the genome, and the loci for which the primer pairs gave the most robust and reproducible amplification (highest PCR efficiency) were kept for experiments.

To place these mispackaging frequencies in perspective - podoviruses (T7-like cyanophages) that infect *Prochlorococcus* have been measured at concentration ranging from 3 to 7 × 10^5^ ml^-1^ in a typical seawater sample from their habitat [20] - which means that the frequencies we observed for podovirus P-SSP7 would extrapolate to a potential of up to ~13 to 30 Mbp of encapsidated host DNA per ml of seawater - roughly a 10^-4^ fraction of total *Prochlorococcus* DNA (see calculation in methods). Thus, even relatively low frequencies of DNA mispackaging inside capsids can represent a significant amount of *Prochlorococcus* genetic information available for horizontal gene transfer in the global ocean at any one time.

### Effect of cell physiology on mispackaging

We next tested if different environmental factors, known to directly affect *Prochlorococcus* physiology, could have an impact on the mispackaging level of the three phages. We first grew the cells at different light intensities, a variable that not only influences the growth rate of *Prochlorococcus* [31] but also influences infection dynamics [32]-[34]. In fact, many cyanophages carry photosynthesis genes in their genomes to boost light-harvesting during infection [35]-[37]; like their hosts, cyanophage genes show day-light cycle transcriptional rhythms [32], [38]; and some cyanophages, including the myovirus P-HM2 used in this study, possess light-specific attachment mechanisms to ensure infection during daylight [32], [39].

Increasing growth light intensity significantly increased the mispackaging frequency – by orders of magnitude – for the podo- and siphovirus but did not impact the myovirus (Fig. 1D). Because the growth rate of the cultures also increases with light intensity [31] it is difficult to separate the effect of light vs growth rate. Examining the mispackaging frequency as a function of growth rate revealed, however, that growth rate alone cannot explain the results (Supplementary Fig. 1); mispackaging was greatest at the highest light intensity that was slightly inhibitory for growth. Thus, we hypothesized that mispackaging in the podo- and siphovirus might be caused by a physiological property related to the growth light intensity. We tested two such physiological parameters: reactive oxygen species toxicity, and the rate of protein synthesis.

### The role of Reactive Oxygen Species in mediating differential levels of mispackaging inside capsids

Reactive Oxygen Species (ROS) are abundant in the surface ocean [40]-[43], and because it lacks the ability to produce catalase, *Prochlorococcus* is highly sensitive and dependent on other species to detoxify them [42], [44]-[47]. The light intensities used in our experiments did, for the most part, result in an increase in intracellular ROS relative to cells kept in the dark (Supplementary Fig. 2). We thus tested different ways of modulating the ROS toxicity in the culture medium and looked at the impact on mispackaging frequency. To increase intracellular ROS levels, we used the herbicide paraquat, which is known to capture electrons from photosystem I resulting in ROS production [48], [49]. A paraquat treatment at 20 μM at the time of infection triggered a significant increase in the level of mispackaging inside P-SSP7 (Fig. 2A). To decrease intracellular ROS levels, we tested a co-culture of *Prochlorococcus* MED4 grown over many generations with the “helper” heterotrophic *Alteromonas macleodii* strain MIT1002 - known to reduce ROS and *Prochlorococcus* oxidative stress via its catalase activity [42], [44], [50] - and compared it with the axenic control. The co-culture showed a dramatic decrease in the level of mispacking inside P-SSP7 capsids (Fig. 2B), supporting our hypothesis that oxidative stress likely plays a role in the frequency of mispackaging.

**Fig. 2.**
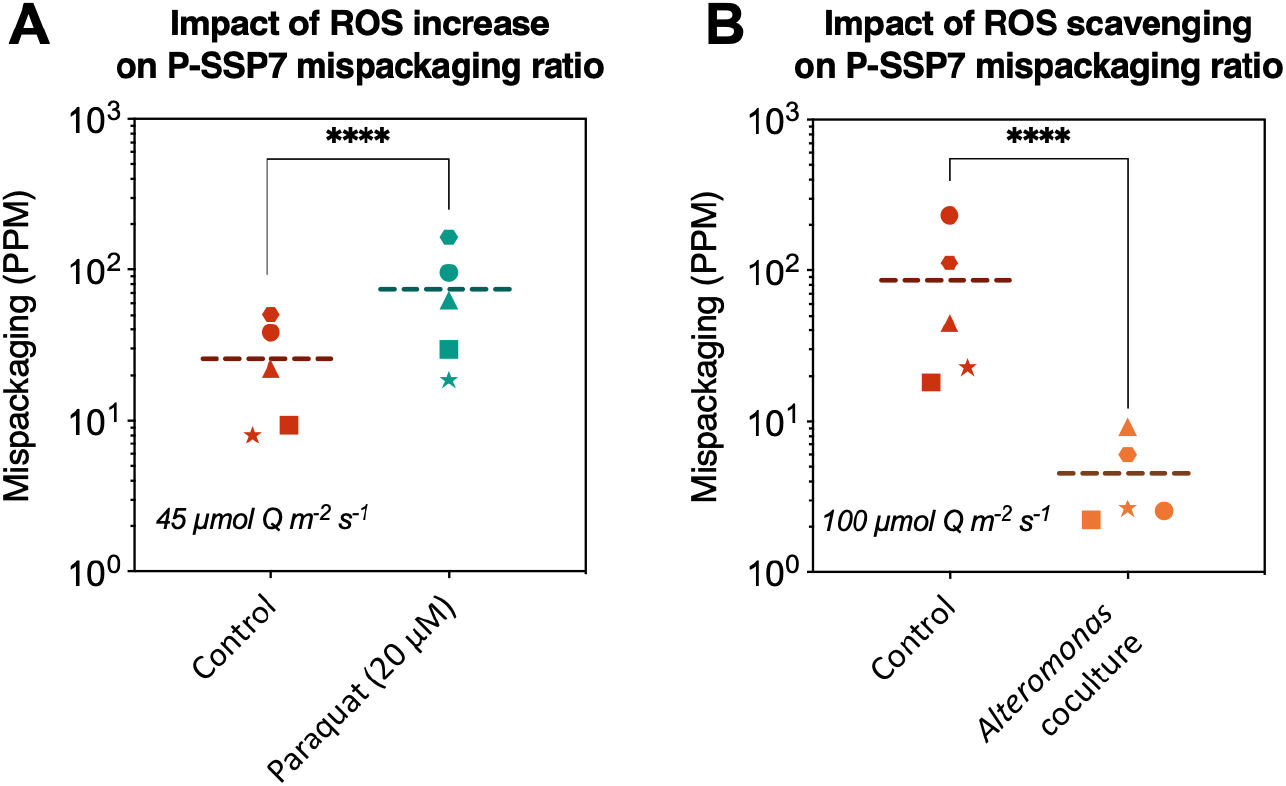
Mispackaging frequency of podovirus P-SSP7 as a function of increased oxidative stress (ROS) induced by Paraquat, and (right) decreased oxidative stress induced by the presence of *Alteromonas,* a “helper bacteria” known to reduce oxidative stress when co-cultured with *Prochlorococcus* [44]. The ROS increase experiments were performed at 45 μmol Q m^-2^ s^-1^, to start at a moderate level of ROS inside cells, and the ROS scavenging experiments at 100 μmol Q m^-2^ s^-1^, to compare the effect with a control at an elevated ROS level.

Overall, the results suggest that ROS could play a role in mispackaging of host DNA inside cyanophage capsids, helping explain the positive correlation between light intensity and the mispackaging level. That the helper strain, *Alteromonas*, could entirely mitigate the impact of high light intensity on mispackaging frequency was somewhat surprising. While at first glance this makes extrapolating our results to field conditions –where *Prochlorococcus* is surrounded by other bacteria – challenging, there are several things to consider. First, *Prochlorococcus* cells in the upper meter of the surface oceans can experience irradiance levels well above the maximum light level used in this study so the ROS detoxification challenges under those conditions could exceed those encountered in our cultures. Second, copiotrophic bacteria like *Alteromonas* are not the dominant bacteria in *Prochlorococcus’* habitat. And finally, other sources of ROS exist in the wild, such as rainfall or the production by other microbes [46], [51]. Thus although mispackaging could be alleviated by the presence of *Alteromonas* in extremely dense laboratory cultures, this does not mean that it does not occur in the wild.

### Accumulation of empty capsids during infection may lead to a higher mispackaging frequency

The rate of protein synthesis is another physiological parameter directly influenced by growth light intensity. In the cyanobacterium *Synechocystis* sp. PCC 6803, for example, there is a general upregulation of the translational machinery with increasing light intensity, which keeps increasing even under conditions of photoinhibition, presumably due to increased turnover of proteins subject to photodamage [52]. Moreover, Puxty et al. recently described that increased light levels during infection of a *Synechococcus* myovirus resulted in faster capsid production, while the phage DNA synthesis remained unchanged [34]. Thus, we hypothesized that if protein translation rates increase to levels that the phage cannot regulate, this could result in an accumulation of empty capsids during infection, increasing the likelihood of host DNA mispackaging events. Indeed, whatever the mechanism by which the host DNA gets loaded inside an empty capsid, the accumulation of latent empty capsids should in turn increase the mispackaging frequency, as long as host DNA is present. Data for marine *Synechococcus* myovirus Syn9 [53], [54] *Prochlorococcus* myovirus P-HM2 [33], [54] and podovirus P-SSP7 [55] shows that host DNA degradation happens quickly, within a few hours of infection, but host DNA is never entirely depleted, suggesting that leftover fragments remain available for mispackaging inside capsids until the cell lyses.

Since DNA damage is a consequence of ROS toxicity [43], [56], ROS serves to augment the imbalance between phage protein synthesis (capsid production) and phage DNA synthesis. Kolowrat et al. showed that long-term acclimation of *Prochlorococcus* to chronic high light and UV exposure leads to a delay in chromosome replication, probably caused by DNA lesions and replication fork arrests which slow DNA synthesis [57]. Thus, whether by increasing protein translation rate (effect of higher light intensity), or decreasing the phage DNA replication efficiency (effect of ROS), this would lead to an imbalance between phage DNA replication and capsid production leading to the accumulation of latent empty capsids waiting for phage genome replication.

We first explored this ‘empty capsid’ hypothesis by inhibiting protein translation and DNA synthesis via the introduction of sublethal doses [58] of the antibiotics chloramphenicol and ciprofloxacin, respectively. Both antibiotics are expected to impact the phage infection but via inhibition of two different host metabolic activities essential for phage particle production: protein translation (chloramphenicol) and DNA synthesis (ciprofloxacin) [59]. We added the antibiotics at the time of infection with the podovirus and observed that, according to our prediction, inhibiting protein translation tended to decrease the frequency of mispackaging while inhibiting DNA synthesis tended to increase it (though the effect was barely significant) (Fig. 3A).

**Fig. 3.**
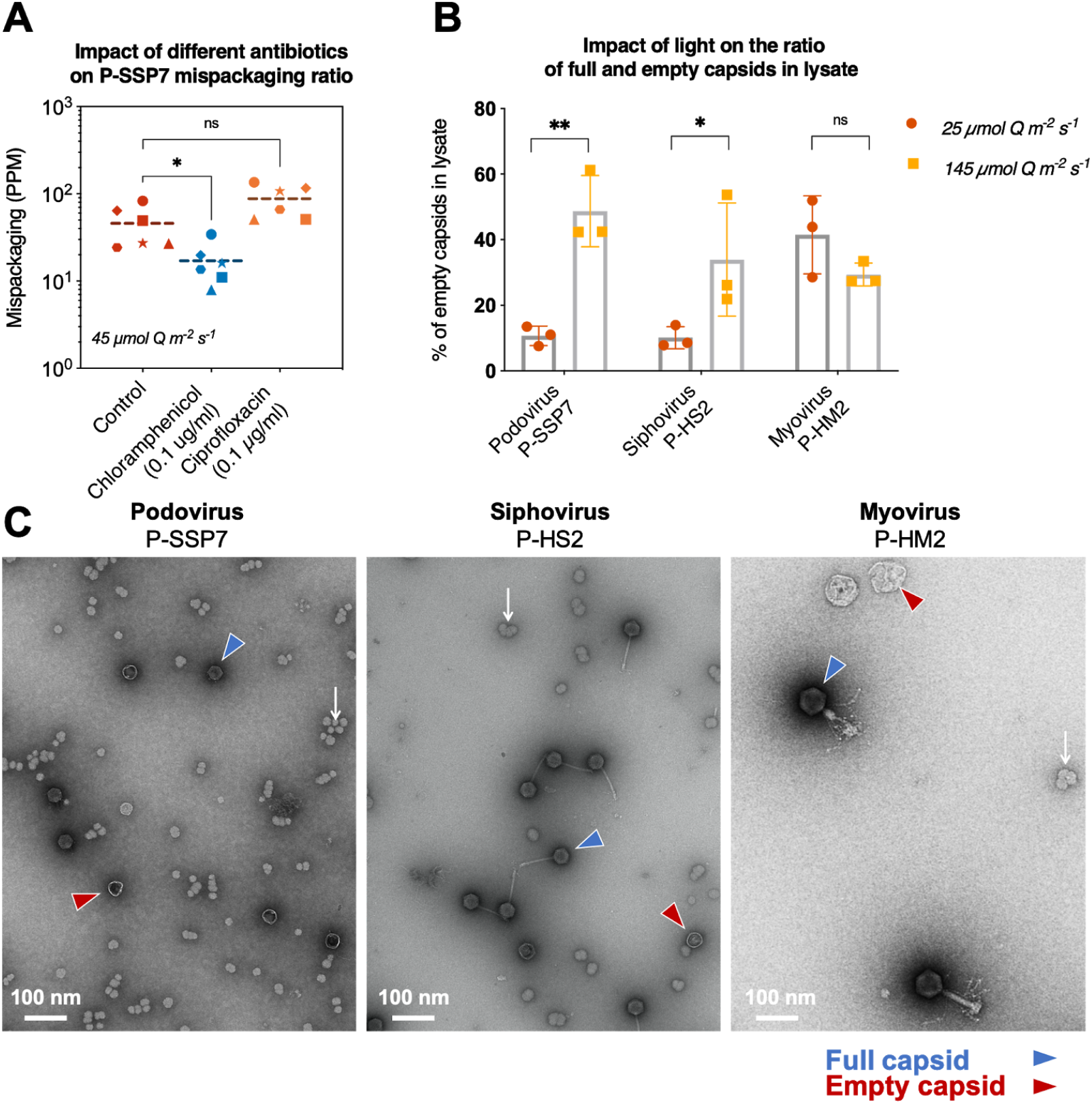
**A)** Impact of protein (chloramphenicol) or DNA synthesis (ciprofloxacin) inhibition on mispackaging frequency during infection of MED4 by the podovirus. The ciprofloxacin increased the overall mispackaging level compared to the control, but slightly below statistical significance (p-value = 0.1091). **B)** Fraction (%) of empty capsids in the lysate at two light levels. **C)** Representative electron micrograph of each cyanophage lysate revealing full (blue arrow) and empty (red arrow) capsids. White arrows show glycogen granules.

As a more direct way to test the hypothesis, we used negative stain transmission electron microscopy to evaluate the proportion of full and empty capsids from the different treatments (Fig. 3B,C, Supplementary Fig. 3,4); more specifically, we measured ratios of full to empty capsids during infection by the three phages at two light intensities (Fig. 3B). The fraction of empty capsids rose from ~10% to ~50% for the podovirus with an increase in light intensity - supporting the idea that empty capsids should accumulate at higher light intensities (Fig. 3B). A similar trend was seen in the siphovirus. In contrast, the fraction of empty capsids for the myovirus did not change significantly as a function of light intensity and was roughly 30% - 40% under both conditions - consistent with lack of light intensity-dependent mispackaging for this phage (Fig 1D).

The different behavior in response to light of the podo and siphovirus compared to the myovirus - both in terms of mispackaging and full/empty capsid ratios - cannot be explained solely from their different DNA packaging mechanisms (Table 2). It could at least partly be explained by auxiliary metabolic genes (AMGs) encoded in their genome (Table 2). The myovirus P-HM2 genome encodes for 11 different AMGs, including a CP12 homolog, which inhibits the Calvin cycle activity, and a TalC homolog, which reroutes resources to the pentose phosphate pathway for nucleotide production [54], while the podovirus P-SSP7 encodes only 4 AMGs including TalC, and the siphovirus P-HS2 encodes none (though it possesses other unknown ways to influence the host metabolism) [60]. Thus, the myovirus - better equipped to ‘control’ the host metabolism - might be more resilient in the face of different metabolic regimes imposed by the light level, for example.

To explore this aspect further, we measured the burst sizes and latent period for the three cyanophages at two different light levels (Table 1). Results are in accordance with our prediction since the myovirus infection parameters did not vary in function of light, while it changed widely for both the podovirus and siphovirus. Comparing the podovirus and siphovirus patterns, however, we see distinct differences. While the latent period decreased for both under higher light, burst size decreased for the podovirus and increased for the siphovirus. This result was unexpected, given their similar responses to light in terms of host DNA mispackaging and the proportion of empty capsids. This suggests that mispackaging is independent of cyanophage fitness.

## SYNTHESIS

This is the first report of the encapsidation of *Prochlorococcus* DNA in the capsids of cyanophages. While the myovirus showed a relatively low mispackaging level regardless of conditions, the podo- and siphoviruses showed significant levels of mispackaging at higher light intensities, representing a potential vector for horizontal gene exchange in *Prochlorococcus* by transduction. Our working model proposes that the dependence of mispackaging on light intensity may involve ROS production and/or an imbalance between DNA and protein synthesis during infection (Fig. 4) - a tendency that the myovirus P-HM2 might be able to mitigate thanks to tighter control of the host metabolism via AMGs in its genome.

**Fig. 4.**
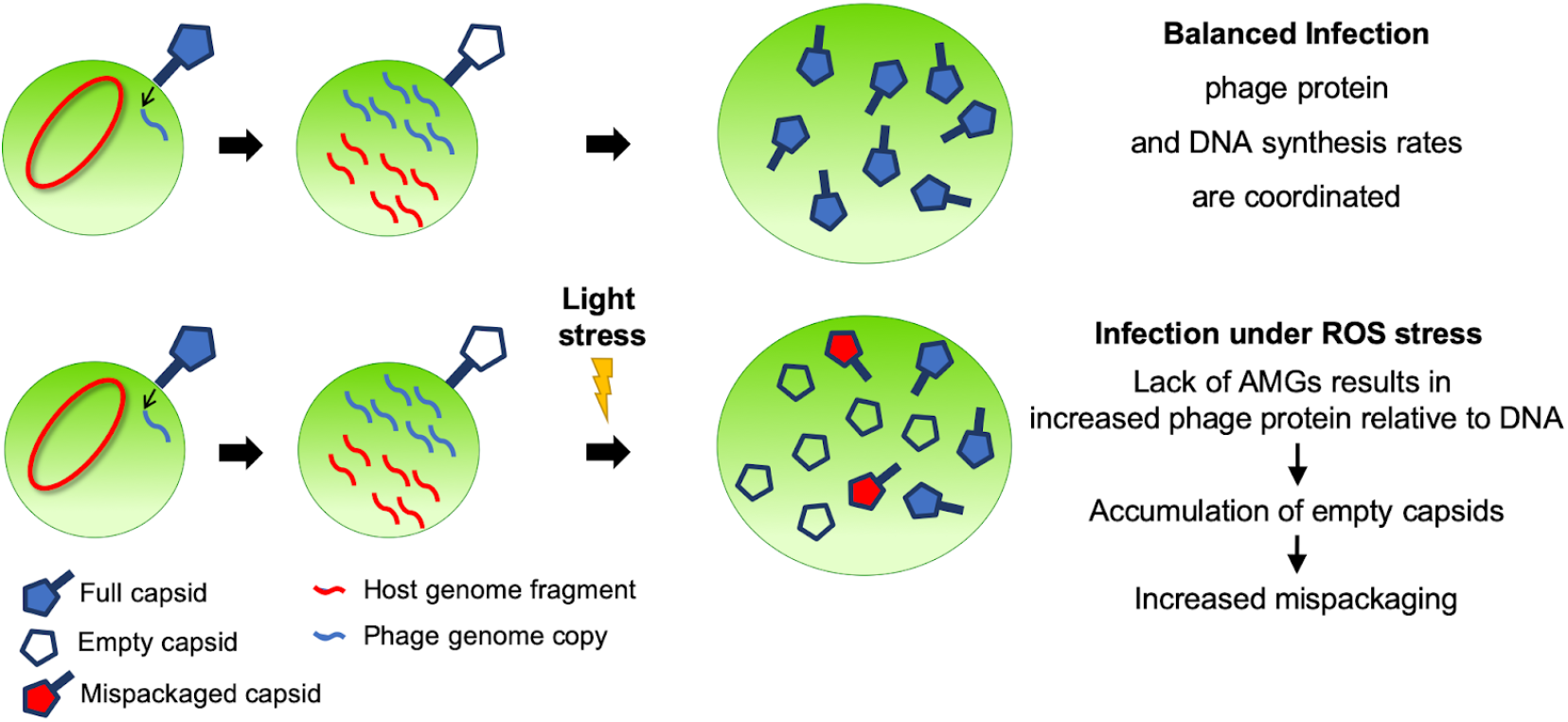
A working model explaining the increased mispackaging of host DNA inside capsids by the podovirus and siphovirus. Higher light levels produce higher intracellular ROS levels and push the host metabolism toward faster protein translation and/or slower DNA synthesis which causes an imbalance in the production rate of capsids and cyanophage genome copies in favor of capsids - a tendency that the myovirus P-HM2 might be able to mitigate thanks to a tighter control of the host metabolism via Auxiliary Metabolic Genes (AMGs) in this cyanophage’s genome. The higher number of latent empty capsids during infection would then increase the likelihood of host DNA mispackaging events.

Interestingly, the underlying mechanisms we describe behind the mispackaging of host DNA inside cyanophage capsids are likely to extend beyond cyanophages. In particular, we observed that mispackaging can be impacted either positively or negatively by antibiotics, a result that could be relevant to the field of phage therapy [61], [62] which is often used in conjunction with antibiotics. In such instances, lateral transfer of DNA through phage capsids could have the unwanted effect of spreading antibiotic resistance genes.

Many unknowns remain to be studied before we can quantify with any certainty the amount of transduction happening among *Prochlorococcus* cells in the marine environment. For example, only a subset cyanophage capsids deliver their content, and when they do, multiple barriers exist to prevent the recombination of the incoming exogenous DNA in the recipient cell [13], [14], [63]. Future directions to advance our understanding of transduction in the marine environment will involve the direct measurement of host DNA encapsidation in environmental samples [64]-[66], or if ways to prepare cyanophage capsids loaded with known and traceable DNA are developed, the direct analysis of transduction events [16].

## MATERIALS AND METHODS

### Culture conditions

Axenic *Prochlorococcus* MED4 cells were grown under constant light flux (45 μmol photons m^-2^ s^-1^ unless otherwise specified) at 24°C in natural seawater-based Pro99 medium containing 0.2-μm-filtered Sargasso Seawater, amended with Pro99 nutrients (N, P, and trace metals) as previously described [67]. Growth was monitored using bulk culture fluorescence measured with a 10AU fluorometer (Turner Designs).

For the coculture, *Alteromonas macleodii* strain MIT1002 [68] (maintained in ProMM medium - Pro99 medium, as above, plus lactate, pyruvate, glycerol, acetate, and Va vitamins [69]) was spun down and washed twice in Pro99 medium to minimize carryover of trace organic compounds prior to being added to the *Prochlorococcus* MED4 cultures. The coculture was acclimated to constant light flux along with axenic cultures used for comparison.

### Encapsidated DNA extraction

We have set up a protocol to purify *Prochlorococcus* cyanophage-encapsidated DNA free of any host DNA contamination, based on published procedures [29], [30], [64]. Briefly, we omitted the cesium chloride gradient step to optimize the DNA yield per sample, and the potential presence of vesicle-encapsulated DNA was removed by chloroform treatment [11].

30 mL cultures of exponentially growing *Prochlorococcus* MED4 cells were infected with cyanophage P-SSP7, P-HS2 or P-HM2 at a multiplicity of infection (MOI) ~0.1. The MOI had no effect on the mispackaging frequency (Supplementary Fig. 5). Cultures were pre-acclimated to the specified light level for several transfers prior to infection. After complete lysis, the lysates were centrifuged at 7,000 g for 20 min and filtered on 0.2 μm Steriflip filters (MilliporeSigma). Lysates were then concentrated using 15 mL Amicon centrifugal concentrators 100 kDa cutoff (MilliporeSigma) down to a volume of ~2 mL. Samples were treated with 10% volume of chloroform, and washed two times in SM buffer (50 mM Tris pH=7.5, 100 mM NaCl, 8 mM MgSO_4_) using a 4 mL Amicon centrifugal concentrators 100 kDa cutoff (MilliporeSigma). Of note, the lysate volume was never concentrated below 0.5 mL during washes, as this resulted in significant losses. As a control for digestion, 0.3 μg of pUC19 plasmid DNA (NEB) was added in the sample. The non-encapsidated DNA was then removed by a 1h incubation at 37°C with 4 U TURBO DNase enzyme (Invitrogen), 1X TURBO DNase Buffer, 0.1 mg mL^-1^, RNase A (Invitrogen) and 1X cOmplete*™* EDTA-free Protease Inhibitor Cocktail (Roche); After 1h, 4 U TURBO DNase enzyme was added and incubated for another hour at 37°C. Finally, the encapsidated DNA was extracted using a standard phenol-chloroform extraction, followed by purification on AMPure XP magnetic beads (Beckman Coulter). Samples were eluted with 50 μL of MilliQ water and used for qPCR quantification.

### Mispackaging quantification by qPCR

DNA samples were quantified using PicoGreen^®^ (ThermoFisher scientific) and quantitative PCR reactions were performed with the QuantiTect Probe PCR Kit (Qiagen), using 1 ng of extracted phage DNA in 25 μL reaction volume. Primers for the 6 amplicons from the *Prochlorococcus* MED4 genome used in all experimental conditions to evaluate mispackaging frequency are listed in Table 3. Amplification was carried out in a CFX96 thermocycler (Bio-Rad), cycling conditions were 15 min at 95 ° C, followed by 40 cycles of denaturing for 15 s at 94 °C and annealing for 30 s at 55 °C. In each run, the DNA sample was compared to a serial dilution of MED4 genomic DNA from 1×10^-9^ g to 1×10^-12^ g. The number of encapsidated gene copies was calculated with the formula:

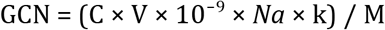

GCN: Gene Copy Number in the PCR reaction; C: Measured concentration from standard curve (ng/μL); V: Volume of PCR reaction (μL); M: Molecular weight of host chromosome (g/mol); *Na*: Avogadro’s number; k: Number of gene repeats in the genome. Results are then expressed in gene copies per million cyanophage capsids (PPM), by dividing the GCN value with the number cyanophage genome copies inside 1 ng of cyanophage DNA, times 1 million. According to qPCR control reactions containing no input template, values below 0.5 PPM could be noise (set as the detection limit).

To confirm the complete removal of non-encapsidated DNA, the same qPCR reaction was performed using primers M13F (gtaaaacgacggccagt) and M13R (caggaaacagctatgac) on each sample. If traces of the pUC19 plasmids were detected, samples were considered contaminated and discarded.

### Extrapolation to the natural environment

To extrapolate our results for the podovirus to the natural environment we began with the following assumptions:

1. The value obtained with P-SSP7 applies to all *Prochlorococcus*-infecting podoviruses, measured at concentrations ranging from 300,000 to 700,000 per mL of seawater [20];
2. ‘Mispackaged’ cyanophage capsids are filled with host DNA to capacity, that is ~45 kb, the average podovirus genome size [71];
3. The entire *Prochlorococcus* chromosome can be mispackaged at the mean value we obtained for the 6 loci we tested. So considering the mean value of 25.97 PPM at 45 μmol Q m^-2^ s^-1^ (a conservative value, given that under laboratory conditions, mispackaging reached an average of 87.75 PPM at 145 μmol Q m^-2^ s^-1^), we extrapolate a mispackaging frequency of *25.97 chromosome copies per million phage capsids in the environment*.

A million phage capsids contain 45 kb × 10^6^ = 45,000 Mbp of DNA so that given the assumptions, 25.97 chromosome copies per million phage capsids results in a ratio of phage:host encapsidated DNA of 45,000 Mbp / (25.97 × *(Prochlorococcus* genome size)) = 1,045. In other words, 1 out of 1,045 capsids are filled with host DNA. Considering 300,000 capsids per mL of seawater, that equates to ~13 Mbp of host encapsidated DNA per mL of seawater. Considering 700,000 capsids, it goes up to ~30 Mbp.

Obviously, this estimation is subject to vast uncertainty, but it serves as an indication of how mispackaging values in PPM can relate to the context of *Prochlorococcus* and its phages in the wild.

### Measurement of intracellular ROS

Intracellular ROS levels were measured via H_2_DCFDA assay (ThermoFisher Scientific #D399), a cell-permeant that is oxidized to the fluorescent DCF form in presence of ROS. Fluorescence per cell was measured on an Influx flow cytometer (Becton Dickinson, Franklin Lakes, NJ, USA); cells were excited with a blue 488 nm laser and analyzed for chlorophyll fluorescence (692/40 nm) and DCF (530/40 nm). Calculations of relative fluorescence per cell were done by normalizing red chlorophyll fluorescence per cell to 2 μm reference fluorescent beads (catalog no. 18604; Polysciences, Warrington, PA, USA) as previously described [72]. All flow cytometry data were analyzed using FlowJo version 10.6.1 (FloJo, LLC, Ashland, OR, USA). Triplicate *Prochlorococcus* MED4 cultures acclimated to the different light levels were harvested by centrifugation and resuspended in fresh Pro99 medium before the addition of reconstituted H2DCFDA to a final concentration of 30 μM. Cells were incubated in the dark at room temperature for 30 min, then exposed to their growth light intensity for 30 min, and samples were immediately run on the flow cytometer.

### One-step method for cyanophages growth curve experiment

Infection dynamics were assayed using qPCR to enumerate both intracellular and extracellular cyanophage genome copy number (GCN) [54], [55]. Triplicate mid-exponential cultures acclimated to high light (145 μmol Q m^-2^ s^-1^) and low light (50 μmol Q m^-2^ s^-1^) were infected at MOI ~1. To ensure the same MOI was used for a given phage at the two light levels, cell abundances were measured prior to infection and the addition of the cyanophage stock was adjusted accordingly. After 1 h, cultures were diluted 1:10 in fresh Pro99 medium, to stop further infection. Samples were taken immediately after dilution of the infected cultures and then at 2, 5, 8, 11, 14, 17, 20, 23, 26, 31 h. 200 μL samples were aliquoted into 96-well MultiScreen-HTS GV filter plate with 0.22 μm filters (Millipore, MSGVN2210) and filtered using a MultiScreen-HTS Vacuum Manifold (Millipore) into 96-well polystyrene microplates. Filtrates were used for qPCR quantification of extracellular cyanophage genome copies, while filters were used for qPCR quantification of intracellular cyanophage genome copies. Quantitative PCR conditions were the same as above, using primers gaacacttccgcccttacct and ctgcaacgaaagggaattgt for P-SSP7; cgtagagaaggtggcagagg and gaccttccgatgttaaattgc for P-HM2; gaattgctccaatcgtcgtt and cagctcgtgaaaacatcgaa for P-HS2.

In each biological replicate, the burst size was calculated as the total number of cyanophages produced during the lytic cycle (extracellular cyanophage GCN at the end of the lytic cycle minus extracellular cyanophage GCN at t_0_) divided by the number of infected cells at t_0_ (intracellular cyanophage GCN at t_0_). Note that cyanophage latent period lengths and burst sizes are difficult to establish and subject to significant uncertainty. Both parameters are better regarded as relative measures that are specific to the methods and conditions we used to obtain them.

### Quantification of full and empty capsids fractions in cyanophage lysates by electron microscopy

Triplicate mid-exponential cultures acclimated to high light (145 μmol Q m^-2^ s^-1^) and low light (25 μmol Q m^-2^ s^-1^) were infected at MOI ~0.1. Once lysis of the culture was completed, samples were centrifuged at 7,000 g for 20 min to remove bacterial debris and filtered onto 0.2 μm Steriflip filter units (Millipore Sigma). 4 mL of each clarified lysate was subsequently ultracentrifuged at 32,000 RPM for 1 h, 4°C in SW 60 Ti rotor (Beckman Coulter) to pellet cyanophage particles, which were resuspended in 50 μL of fresh and filtered Pro99 medium. 10 μL drops of each suspension were then placed directly on glow discharged carbon-coated grids (EMS, USA) for 1 minute. The grids were then blot-dried on filter paper, washed on a drop of ultrapure water, and negatively stained with 2% uranyl acetate in water. Specimens were examined on an FEI Tecnai T12 electron microscope operating at 80 kV at nominal magnifications of 18500–48000 and 1–3 μm defocus. Of note, P-HS2 particles were significantly aggregated after resuspension. To allow counting of this phage, Triton X-100 detergent (Sigma-Aldrich) was added to a final concentration of 0.1% for 1 h at room temperature before grid preparation.

The morphology of empty capsids (Fig. 3C, Supplementary Fig. 3) was consistent with examples from the literature [73]-[76], and could easily be differentiated from extracellular membrane vesicles (Supplementary Fig. 4). Empty capsids for the tailed cyanophages P-HM2 and P-HS2 were not tailed, consistent with the fact that they are immature capsids that have not been filled, and not capsids that delivered their content inside cells. A minimum of 100 particles (more frequently 200 to 400 particles) was counted on electron micrographs for each biological replicate to assess the full:empty capsid ratio in the lysate.

## ACKNOWLEDGMENTS

This study was supported in part by the Simons Foundation (Life Sciences Project Award IDs 337262, 509034SCFY17, 647135 and SCOPE Award ID 329108, to SWC), and an NSF-EDGE grant (1645061 to SWC).

We thank Nicki Watson for her help setting up the protocol for TEM visualization of cyanophage capsids, Cameron Haase-Pettingell for her recommendations on how to concentrate and purify cyanophage particles, Debbie Lindell and Dror Shitrit for their recommendations on the best methods to measure cyanophage infection parameters, and Thomas Hackl for his help with mispackaging frequency calculations.

