## Supplementary data for "Frequency of mispackaging of *Prochlorococcus* DNA by cyanophage"

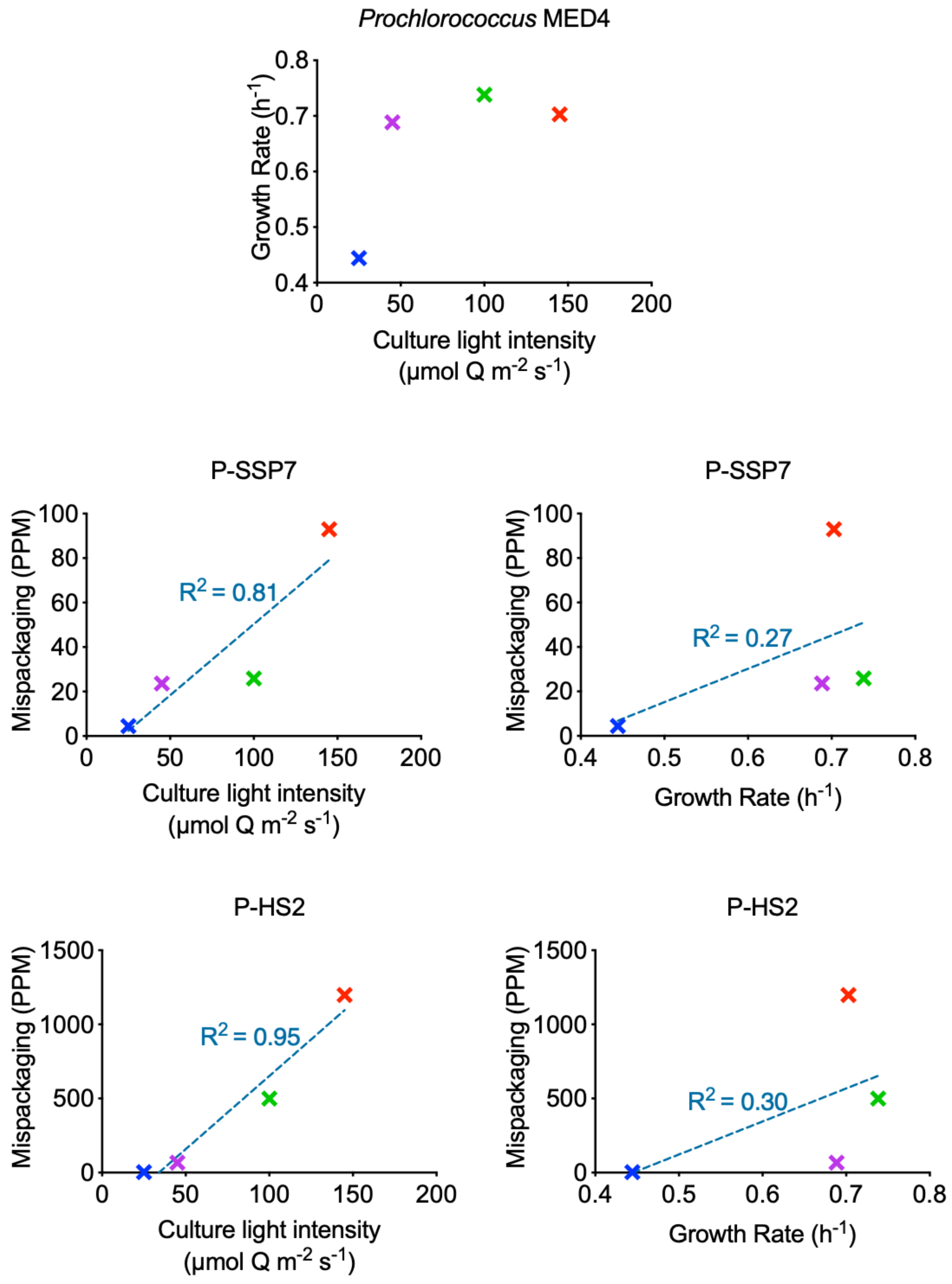


**Supplementary Fig. 1.** Comparison of mispackaging frequency as a function of light intensity or growth rate of the culture. The data points are colored according to the light intensity (see top graph). The blue dotted line corresponds to a linear regression and its corresponding $R^2$ value is displayed. Cells growing at 145 μmol Q $m^{-2}$ $s^{-1}$ (red data points) are photo-inhibited (light stress phase), but the mispackaging level keeps increasing at this high light level, yielding a better correlation to light intensity.

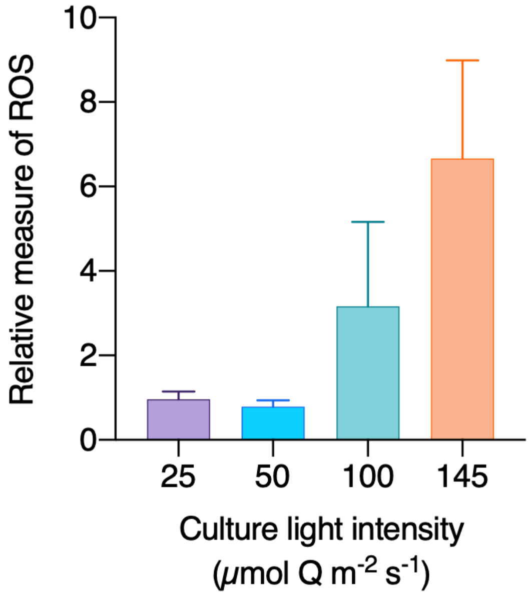


**Supplementary Fig. 2**. Reactive oxygen species (ROS) as a function of growth light intensity of the cultures of *Prochlorococcus.* ROS is displayed as mean DCF (see methods) fluorescence per cell (a measure of intracellular ROS) after a 30 min light treatment, normalized to cells kept in the dark. Light-induced ROS were not detected for 25 and 50 μmol Q $m^{-2}$ $s^{-1}$, but increased gradually for 100 and 145 μmol Q $m^{-2}$ $s^{-1}$.

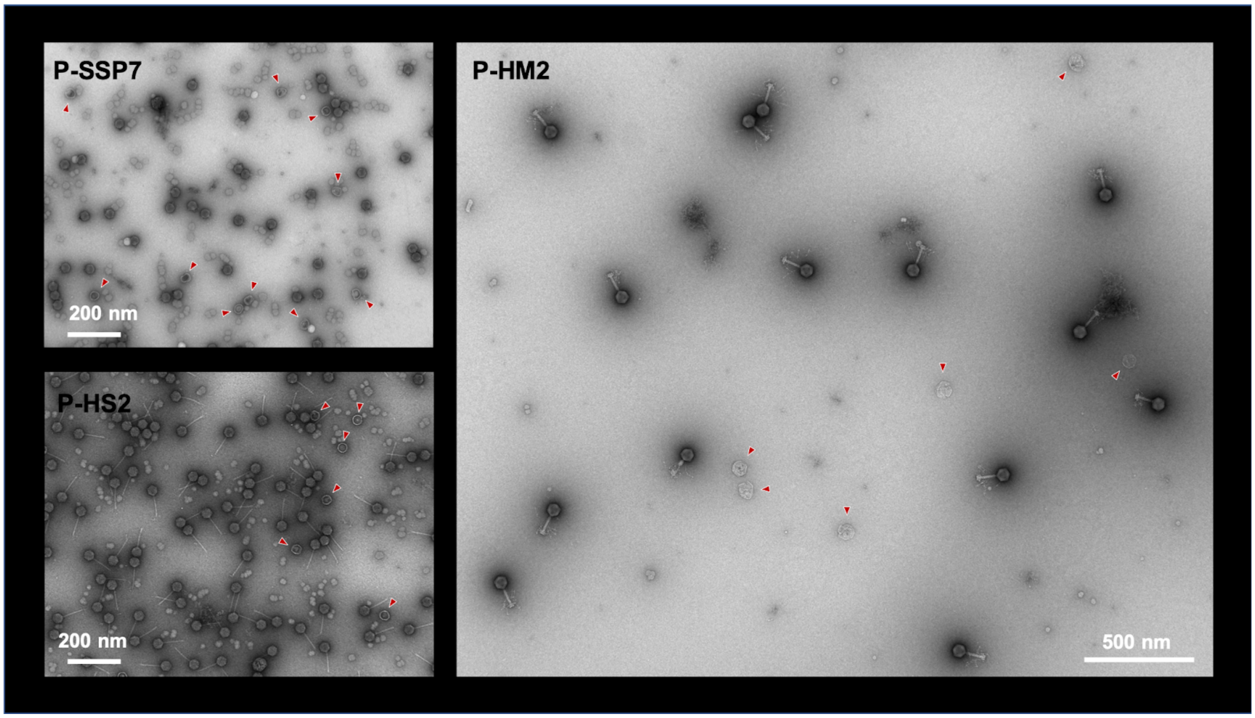


**Supplementary Fig. 3**. Representative electron micrograph of each cyanophage lysate, showing a broader field than Fig. 3C. Empty capsids are indicated with red arrowheads.

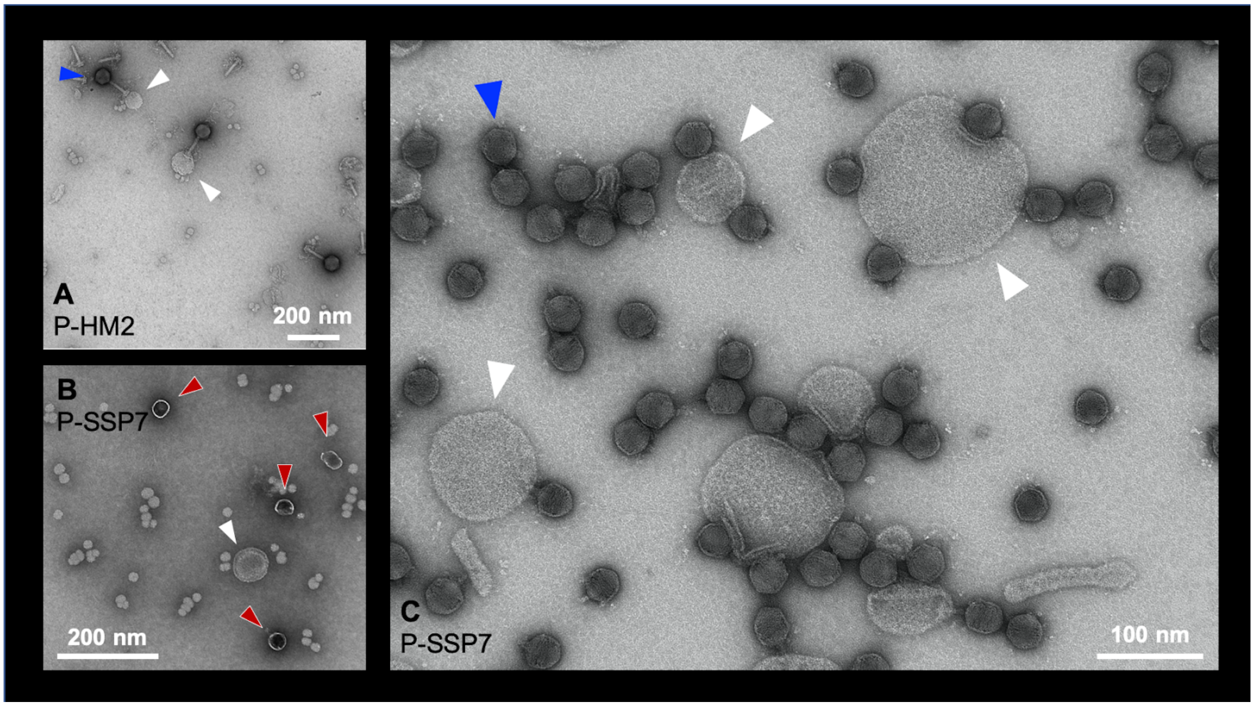


**Supplementary Fig. 4.** Negative stain electron micrographs showing the visual difference between *Prochlorococcus* membrane vesicles (white arrowhead) with empty cyanophage capsids (red arrowheads) and full cyanophage capsids (blue arrowheads). Vesicles appear less contrasted than empty capsids and vary widely in size and shapes. Interestingly, cyanophages were sometimes attached to vesicles (panel A, C). While this could be an artifact of the preparation, it is worth mentioning that this has been observed before [1] and one of the hypotheses for the function of vesicles is that they could serve as decoys for phage infection, reducing phage fitness.

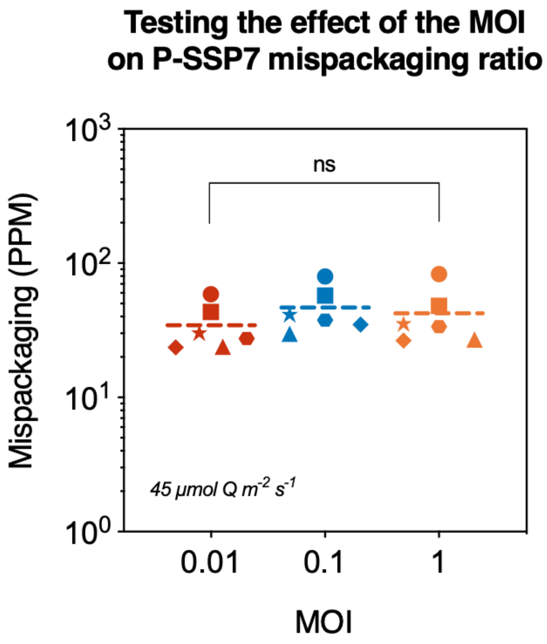


**Supplementary Fig. 5**. Mispackaging frequency as a function of multiplicity of infection for the podovirus P-SSP7 infecting *Prochlorococcus* MED4.
